# A frameshift mutation in the methyltransferase *rlmN* is associated with increased linezolid resistance in *Mycobacterium tuberculosis*

**DOI:** 10.1101/2025.11.20.689544

**Authors:** Bryn Marie Reimer, Anna G. Green

## Abstract

Linezolid is a key component of treatment regimens for multidrug-resistant and extensively drug-resistant tuberculosis, which is caused by the pathogen *Mycobacterium tuberculosis* (MTB). Resistance to linezolid in MTB has traditionally been attributed to mutations in the 23S rRNA (*rrl*) and ribosomal protein L3 (*rplC*), but only a fraction of clinically observed linezolid resistance is explained by mutations in these two genes. We report that an analysis of strains with paired whole-genome sequencing and linezolid minimum inhibitory concentration (MIC) phenotyping from the Bacterial and Viral Bioinformatics Resource Center (BV-BRC) reveals that a relatively common frameshift mutation in MTB methyltransferase *rlmN* (5.3% of assessed isolates) is significantly associated with increased linezolid MIC. In additional to statistical associations, we provide evolutionary evidence of homology to an established linezolid resistance mechanism in *Staphylococcus aureus*, and structural evidence that the frameshift mutation likely ablates *rlmN* methyltransferase functionality.

## Introduction

Tuberculosis (TB) remains the world’s most deadly infectious disease, causing an estimated 1.25 million deaths in 2023 alone.^1^ TB is caused by the bacillus *Mycobacterium tuberculosis* (MTB), which is uniquely difficult to treat with antimicrobial agents, due in part to its waxy mycolic acid coating.^2^ Further complicating treatment is the high rate of drug-resistant infections: in 2023, an estimated 400,000 individuals developed multidrug-resistant or rifampicin-resistant TB, which requires treatment with second-line drugs.^1^

Linezolid, a first-generation oxazolidinone, is a commonly used drug in multidrug- and extensively drug-resistant TB,^3^ and resistance to linezolid is present in an estimated 4.2% of patients with multi-drug resistant TB.^4^ The World Health Organization (WHO) maintains a catalogue of mutations known to cause resistance to antitubercular drugs, including linezolid, based on stringent statistical criteria applied to mutations in candidate genes.^5^ While mutations in *rrl*, the 23S rRNA, and *rplC*, ribosomal protein L3, have been observed in linezolid-resistant strains of MTB,^6,7^ it is unclear what proportion of resistance is explained by these mutations. Estimates vary, but between 20% and 54% of linezolid-resistant isolates appear to have wild-type *rrl* and *rplC*.^8,9^ Because a large proportion of linezolid resistance cannot be easily tied to mutations in these two major candidate genes, it is reasonable to presume other genetic mechanisms of resistance may be present.

In this work, we demonstrate that the presence of a frameshift mutation in the ribosomal methyltransferase *rlmN* is associated with increased quantitative linezolid resistance in MTB as measured by MIC (referred to as “linezolid resistance” hereafter). We analyze MTB isolates with paired genotype and phenotype data to show that the presence of this frameshift mutation is significantly predictive of linezolid resistance, explaining an additional 32.1% of resistant isolates compared to using only known resistance-associated mutations from the WHO.^5^ Additionally, we contribute a brief analysis of RNA methyltransferases in MTB to justify our focus on *rlmN*, which is known to cause antibiotic resistance in the pathogen *Staphylococcus aureus*.^10^ Finally, we contribute a structural analysis of *rlmN* that provides evidence that the observed frameshift mutation would have a deleterious effect.

## Results

### A protein similarity analysis uncovers MTB’s sole *rlmN* homolog

Multiple previous studies have implicated RNA methyltransferase genes in linezolid resistance in other bacterial species. Prior work has proposed a role for the *Staphylococcus aureus* methyltransferase *rlmN* in creating the correct methylation state of the rRNA to facilitate linezolid binding.^10^ In addition, the presence of the plasmid-borne RNA methyltransferase *cfr* in several species such as *S. aureus* and *Escherichia coli* has been associated with linezolid resistance.^11^ The importance of these two methyltransferase genes in linezolid resistance indicates that the correct methylation state of the ribosomal RNA is critical for linezolid function. Therefore, we hypothesize that methyltransferases provide a promising reservoir of genes that may be implicated in linezolid resistance in MTB.

The only prior catalogue of all MTB methyltransferases, published in 2016, does not capture some important members of the MTB RNA methyltransferases (see Discussion).^12^ We therefore contribute a systematic catalogue of RNA methyltransferases with two goals: (1) to provide a candidate gene list for further exploration of drug targets in MTB and (2) to clarify the identity of *rlmN* in MTB, as there has been some confusion in the literature about the identity of *rlmN* in MTB; some have proposed that *tsnR* is its homolog in MTB.^13^ Because a loss of function mutation in the *S. aureus rlmN* leads to increased linezolid resistance,^10^ we hypothesized that if MTB has an *rlmN* homolog, its function may also affect linezolid’s activity. Thus, we proceeded to catalogue the RNA methyltransferases in MTB and to align the sequences of these methyltransferases with the *S. aureus rlmN* to find its homolog(s) in MTB.

First, we performed a search for all annotated genes that are known or suspected RNA methyltransferases in MTB using a combination of UniProt^14^ and several reference genome annotations (see Methods).^15,16^ We identified 19 putative or confirmed RNA methyltransferases in MTB, representing 20 genes in the H37Rv reference genome (see Table 1). All 19 genes were found in UniProt, though not all were reviewed entries. The *rlmN* gene as annotated by UniProt is a fusion of Rv2879c and Rv2880c, which are annotated as separate, overlapping genes in the H37Rv reference genome. We confirmed with a BLAST^17^ search that there were no additional homologs to *S. aureus rlmN* in the MTB genome (see Methods) that were not recovered by our prior searches through genome annotations.

**Table 1.**
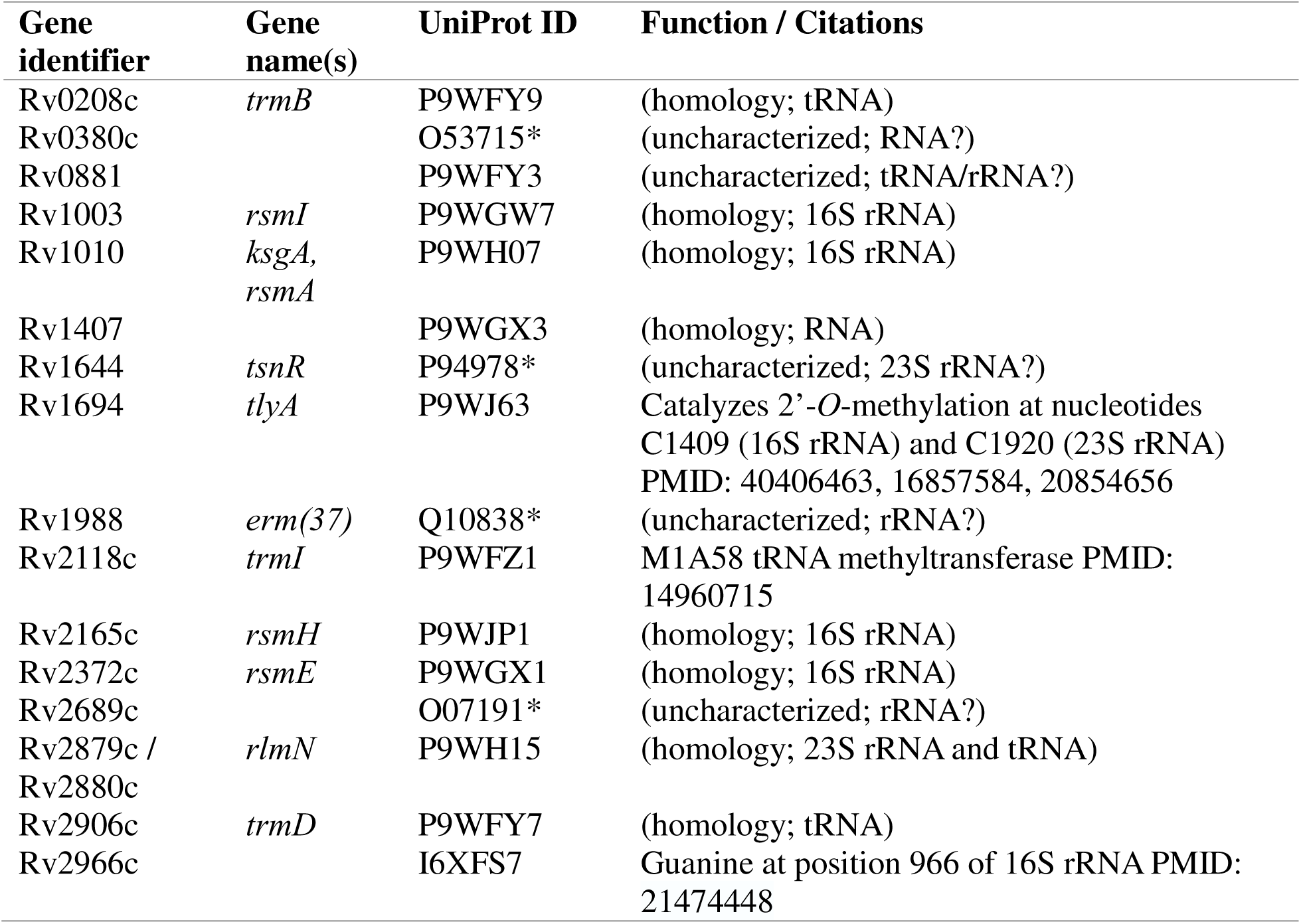
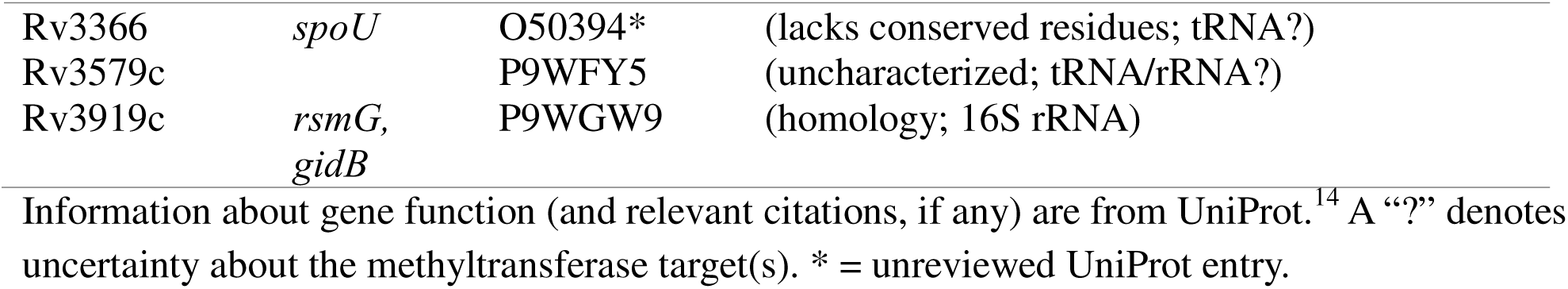
A catalogue of known and suspected RNA methyltransferases in MTB.

We aligned all putative and confirmed RNA methyltransferase protein sequences in MTB to the *S. aureus rlmN* protein sequence (UniProt ID: Q2FZ66) to find potential MTB homolog(s) (see Figure 1). Of all putative and confirmed RNA methyltransferases in MTB, the *S. aureus rlmN* is most similar by percent identity (32.73%) to the MTB *rlmN*, whereas it is only 9.43% identical to MTB *tsnR*. It is a common, if conservative, heuristic to assume homology between two proteins that have at least 30% sequence identity.^18^ Applying this threshold, and noting that BLAST did not find any other potential hits for *rlmN* in the MTB genome, we can confirm that MTB *rlmN* is the sole MTB ortholog to *S. aureus rlmN*. We therefore assert that *tsnR* is not the MTB *rlmN*, nor is it a close homolog of the MTB or *S. aureus rlmN*, despite its similar putative methyltransferase activity, and despite the *in vitro* evidence of *tsnR* association with MTB linezolid resistance.^19^

**Figure 1:**
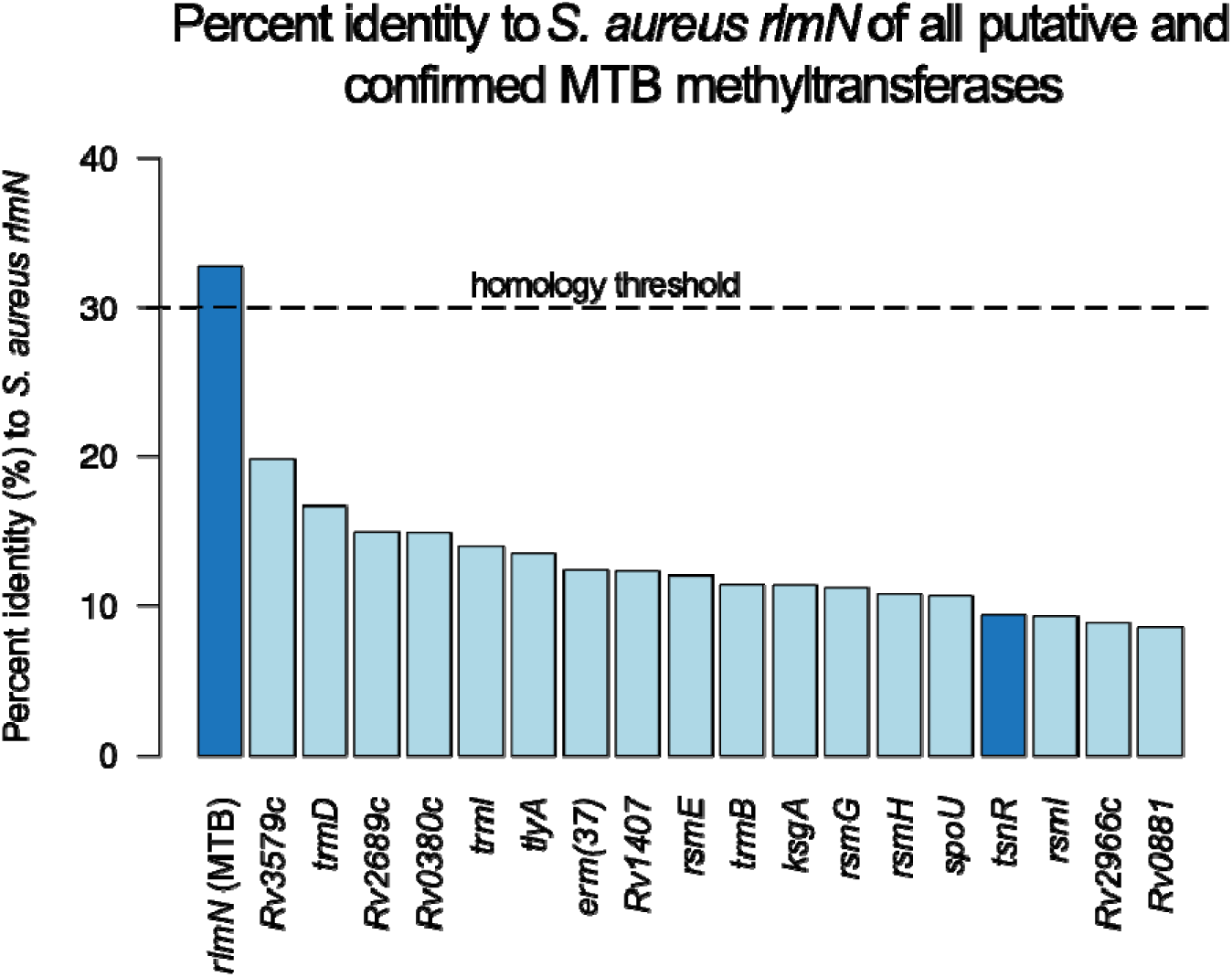
Each putative and confirmed MTB methyltransferase is shown with its associated percent identity to the *S. aureus rlmN*. Highlighted in dark blue are the MTB *rlmN* as well as *tsnR*, which prior literature suggests may be an *rlmN* homolog in MTB. Only the MTB *rlmN* passes the 30% homology threshold to be considered a true homolog.

### Genotypic analysis recovers known resistance mechanisms and reveals that an *rlmN* frameshift mutation is associated with linezolid resistance

We evaluated the association between genomic variants and linezolid resistance by finding genomic variants in sequenced isolates and assessing their association with a change in the observed minimum inhibitory concentration (MIC) for linezolid. By assessing resistance association using MIC values rather than resistance breakpoints, we may be able to find low-level determinants of resistance. Using the BV-BRC as a resource,^20^ we accessed isolates that had (1) lab-determined linezolid resistance phenotypes and (2) a whole-genome sequencing accession number. The genome assemblies associated with the accession numbers for the resulting 8454 MTB isolates were downloaded from AllTheBacteria,^21^ where possible. After pre-processing to remove low-quality genome assemblies or isolates with missing data (see Methods), 7436 isolates remained for analysis.

We used a candidate gene approach for our genotypic analysis, following approach used to generate a WHO-endorsed catalogue of MTB mutations and whether they cause antibiotic resistance (the “WHO mutation catalogue”).^5^ When assessing genotypic variants associated with linezolid resistance, the WHO assess three genes: *rrl*, *rplC*, and *tsnR*, finding mutations significantly associated with resistance in only *rplC* and *rrl* (*rplC* C154R; various in *rrl*, including G2270T). We include these three genes and add *rlmN*. We restricted our search to missense and nonsense mutations for proteins *rplC*, *rlmN,* and *tsnR* and to single nucleotide polymorphisms for the ribosomal genes *rrl*. Indels and frameshifts were included only if they were present in greater than 1% of resistant or sensitive isolates, to minimize possible erroneous mutation calls (see Methods).

For each mutation found in these genes in our dataset, we used the Mann-Whitney U-test to test the null hypothesis that isolates with and without a given mutation are selected from populations with the same MIC distribution (see Methods). We find that six mutations are significantly associated with an increase in linezolid MIC in our dataset (see Table 2). No mutations in *tsnR* attained significant association with resistance in our dataset. Of the six significantly-associated mutations, *rrl* G2270T and *rplC* C154R are also associated with resistance in the WHO mutation catalogue, which uses binarized phenotypes based on the WHO linezolid breakpoint rather than correlations with an increase in MIC.^5^ In addition to these two known variants, we also describe two other *rrl* variants (C344T, C637G) that are significantly associated with resistance in our dataset, but which did not meet criteria for association with resistance in the dataset used by the WHO.

**Table 2:** Observed mutations significantly associated with increased linezolid resistance.

| Gene | Mutation | ΔMIC | MIC<br>+mutation | MIC<br>-mutation | *p*-value<br>(corr.) |
| --- | --- | --- | --- | --- | --- |
| *rrl* | **G2270T** | 2.22 | 2.69 | 0.47 | 2.14e-03 |
| *rplC* | **C154R** | 2.05 | 2.52 | 0.47 | 4.87e-07 |
| *rlmN* | **M176fs** | 0.18 | 0.65 | 0.47 | 2.43e-30 |
| *rrl* | **C344T** | 0.17 | 0.64 | 0.47 | 6.65e-19 |
| *rlmN* * | **D244N\*** | 0.17 | 0.64 | 0.47 | 7.53e-05 |
| *rrl* | **C637G** | 0.17 | 0.64 | 0.47 | 4.13e-05 |

This table reports all assessed mutations which are significantly associated with increased linezolid resistance (Bonferroni-corrected *p*-values < 0.05, $\Delta$ MIC > 0). Also shown is the mean MIC for isolates with the relevant mutation (“MIC +mutation”) and without the mutation (“MIC -mutation”). All *p*-values were calculated using the two-sided Mann-Whitney U-test, and corrected using the Bonferroni procedure. \* = this mutation is only present on the background of the frameshift mutation; see main text.

The final two observed mutations that are associated with resistance in our dataset are both in *rlmN*. The first is an aspartic acid to asparagine missense mutation at amino acid position 244 in the *rlmN* gene (D244N). The second is a frameshift mutation starting at amino acid position 176. This frameshift mutation is relatively common in our dataset (394/7436, or approximately 5.3%). When examining raw MIC values, we find that the presence of this frameshift mutation in any isolate is significantly associated with a raw difference of 0.18 mg/L in MIC (Table 2). If we binarize the phenotypes using the WHO breakpoint of 1.0 mg/L^22^ (isolates with MIC > 1.0 mg/L are resistant), we observe that this mutation is more common in linezolid-resistant isolates (11/128, or approximately 8.6%) than in linezolid-susceptible isolates (383/7308, or approximately 5.2%).

The *rlmN* D244N mutation appears to be significantly associated with resistance in our dataset, but further analysis shows that this mutation is only present on the background of the frameshift mutation. Because D244N mutation appears only downstream of the frameshift mutation at position 176, it is difficult to assess its importance in this dataset. D244N seems likely to be acting as a marker of the frameshift mutation being present.

We also examined how many resistant isolates harbor the mutations that we identify as having significant associations with linezolid resistance (see Table 2). Of the 128 resistant isolates in our dataset (as calculated using the WHO breakpoint), only 28 (21.88%) were found to have at least one mutation from the WHO catalogue of resistance-associated mutations. When we added just the *rlmN* frameshift mutation to this analysis, 37 (28.91%) isolates harbored the frameshift mutation or at least one mutation from the WHO catalogue. Therefore, the frameshift mutation in *rlmN* provides increased explanatory power (32.1%) when assessing the known genetic causes of linezolid resistance, although our baseline ability to ascribe resistance to genetic loci is still limited.

### The frameshift mutation will likely cause loss of function of MTB *rlmN*

To evaluate a possible mechanistic explanation for the observed association, we performed *in silico* protein structure analyses to assess the likely effect of the frameshift mutation on protein function. While there are no experimentally-derived protein structures available for either MTB RlmN or *S. aureus* RlmN, there are several X-ray structures available for *E. coli* RlmN. We aligned the MTB *rlmN* protein sequence with the *E. coli rlmN* protein sequence and found 37% sequence identity across the sequence match, supporting the homology of these two proteins (see Figure 2A). We then generated a protein-ligand interaction fingerprint (PLIF; see Methods) to assess which amino acids were likely important for binding *S*-adenosyl methionine (SAM), the critical cofactor for RlmN protein methyltransferase activity (see Figure 2B). The PLIF shows 5 residues with critical binding interactions with SAM, four of which are ablated when the frameshift mutation is present. Although the frameshift mutation only occurs about halfway through the protein, it clearly impacts important residues for cofactor binding and therefore activity (see Figure 2C), leading us to hypothesize that the frameshift mutation will likely cause loss of function of the MTB RlmN. This loss of function, and consequent loss of rRNA methylation, may be the mechanism of the observed associated linezolid resistance, and it is consistent with the mechanism of resistance observed in *S. aureus*, where abolishment of RlmN activity leads to resistance due to a lack of A2503 methylation.^10^

**Figure 2.**
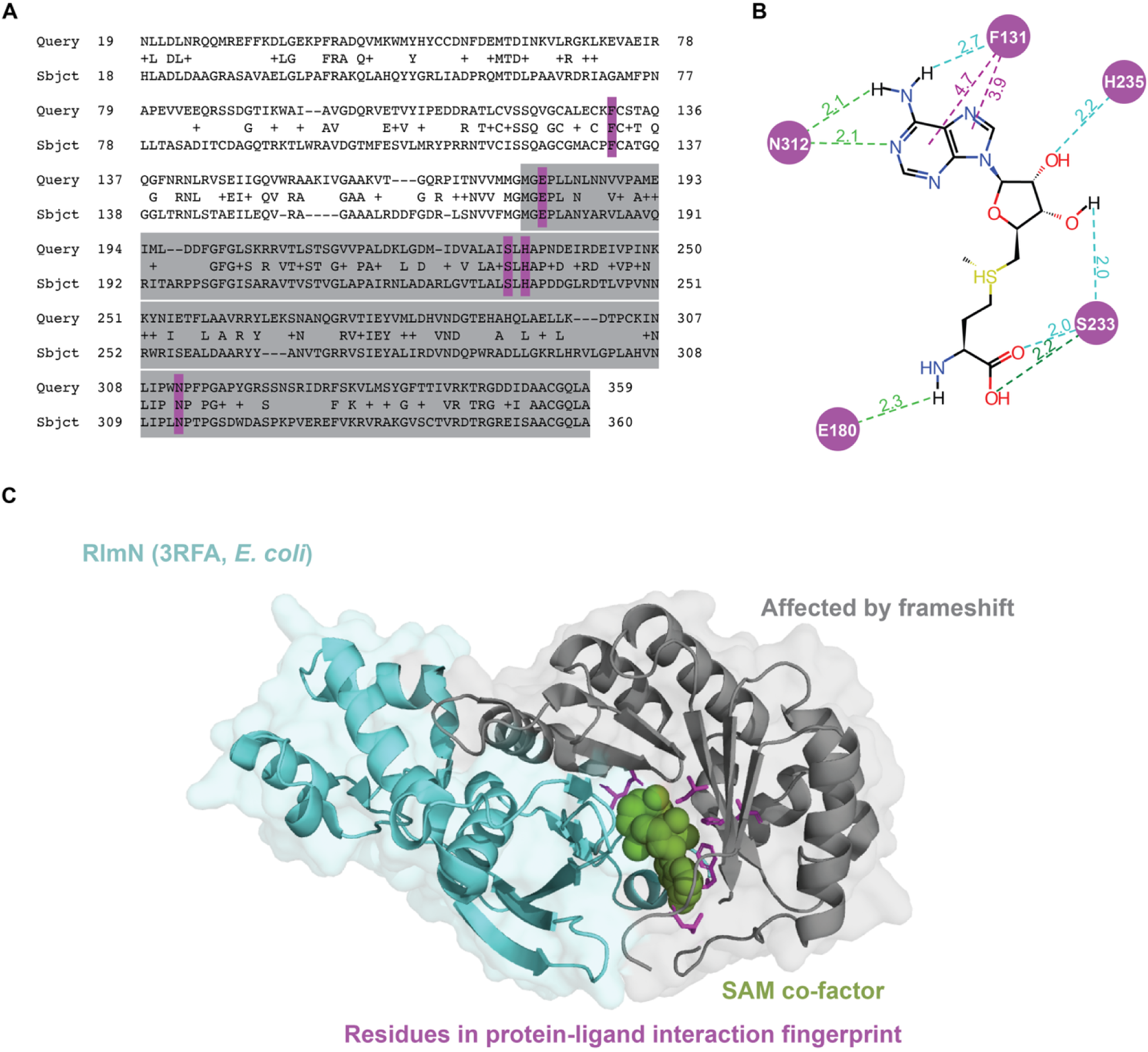
(A) This panel shows the BLAST alignment of the protein sequences for *rlmN* in both *E. coli* and MTB, with grey highlights showing the parts of the sequence that would be downstream of the frameshift mutation. Purple highlights indicate important residues in the protein-ligand interaction fingerprint (PLIF). (B) The PLIF, as calculated by the Cresset software (see Methods), is shown for *E. coli* RlmN (PDBID: 3RFA). The green and blue dotted lines indicate hydrogen bonds and weak hydrogen bonds, respectively. The purple lines indicate aromatic interactions. Amino acids are numbered by their index in *E. coli*. Image initially generated using Flare™ from Cresset®, then modified by the authors. (C) The *E. coli* RlmN protein structure is shown, with the area downstream of the frameshift in grey, the SAM co-factor in green, and residues important in the PLIF in purple.

## Discussion

In this study we report the existence of a significant association between a frameshift mutation in an MTB methyltransferase, *rlmN*, and phenotypic resistance to linezolid as quantified by MIC. To support our addition of *rlmN* to a small list of candidate genes, we contribute a catalogue of all known and putative MTB RNA methyltransferases (Table 1). We demonstrate the homology of MTB *rlmN* to *S. aureus rlmN*, dispelling prior suggestions that *tsnR* may be functioning as an *rlmN* homolog. Upon finding a significant association between a frameshift mutation in *rlmN* and an increase in linezolid MIC, we provide structural evidence for a hypothesized mechanism of resistance.

Despite our survey of all known associated genetic loci for linezolid resistance, using our six reported significantly-associated mutations as well as all WHO-reported mutations that were also observed in our dataset, we were only able to assign a putative genetic cause of resistance to 37/128 (28.91%) of resistant isolates in our dataset. In contrast, Kadura *et al* report that, across eight studies with sequencing results for *rrl* and *rplC*, 85/106 (80.19%) of resistant isolates harbored at least one mutation in those genes.^9^ If we were to use these same criteria and assess isolates with any mutation in *rrl* or *rplC* in our dataset, we would nonetheless only mechanistically explain 42/128 (32.81%) of resistant isolates in our dataset. Notably, in our study, 977/7308 (13.37%) of *susceptible* isolates still had at least one mutation in *rrl* or *rplC*, but that number drops to 396/7308 (5.42%) when only considering significantly-resistance-associated *rrl* or *rplC* mutations, further justifying our choice to restrict analysis to mutations that are significantly associated with resistance.

Regardless, there is a substantial portion of linezolid-resistant MTB isolates where resistance cannot be ascribed to one of the known genetic loci, including our newly-discovered *rlmN* frameshift mutation. This phenomenon of unexplained resistance has also been noted in *Mycobacteroides abscessus*, a closely related mycobacterial species that also only harbors one copy of ribosomal DNA,^23^ where it has been reported that only 8.2% of linezolid-resistant strains contained ribosomal mutations.^24^ A recent review proposes that that mechanisms involving efflux pumps^24^ and the fatty acyl-AMP ligase *fadD32*^25^ may contribute to part of the remaining unexplained resistance.^13^ Future work assessing the importance of mutations in the MTB homologues of these genes could aid in our understanding of currently-unexplained linezolid resistance.

Additionally, while the present work focused on a single methyltransferase, *rlmN*, that has compelling evidence for conferring linezolid resistance in *S. aureus*, many other MTB RNA methyltransferases may also be involved in creating the correct methylation state for linezolid and other ribosome-targeted therapeutics. There has been growing interest in the plausibility of methyltransferases as drug targets for MTB,^26^ in no small part due to the importance of MTB methyltransferases such as *tlyA* in capreomycin resistance^27^ and *gidB* in streptomycin resistance.^28^ We recommend that relevant ribosomal methyltransferase genes be added to candidate gene lists for assessing genotype-phenotype correlations in drug-resistant TB, especially for drugs which target the ribosome. However, we recognize that because the precise mechanism of action is only known for a few of these methyltransferases, picking a targeted candidate gene list based on the proclivity of a gene to methylate near a drug binding site remains difficult. It is plausible that other RNA modifications, such as formylation, acetylation, or sulfation may also be important for drug resistance. Knowledge of the precise RNA modifications required for drug activity can help shape structure-activity studies as we continue to develop ribosome-targeted drugs in the future.

To this end, one of our contributions is a systematic catalogue of all known or putative MTB RNA methyltransferases. We note that a 2016 review of MTB methyltransferases by Grover *et al*. suggests the presence of only 16 RNA methyltransferases, in contrast to our 19 identified RNA methyltransferases.^12^ Their list of methyltransferases with RNA-binding activity does not recognize the RNA binding potential of Rv1407, Rv2689c, Rv3919c, and Rv2966c, which our list does due to their homology to known methyltransferases in other species. Rv2966c, in particular, is now an experimentally-verified MTB rRNA methyltransferase.^29^ Additionally, Grover et al consider Rv3024c a methyltransferase; however, more recent evidence from 2023 supports the role of Rv3024c as an enzyme that sulfurates (rather than methylates) tRNA isoacceptors.^30^

In our study we include two methyltransferases as candidate genes for association with linezolid resistance: *rlmN*, our gene of interest, and *tsnR*, which is on the current candidate gene list for linezolid resistance in the WHO mutation catalogue.^5^ Loss of function of *tsnR* has been observed *in vivo* to correlate to linezolid resistance.^19^ Despite this observation *in vitro*, no observed *tsnR* mutations met our stringent thresholds for significant association with resistance. This result was also seen in the WHO mutation catalogue, which reports no *tsnR* mutations as being associated with resistance.

The association of *rlmN* mutation with linezolid resistance in MTB has not been reported previously. While Li et al performed a genome-wide CRISPRi screen that covered 98% of the MTB genome, neither *rlmN* nor its two constituent annotated genes in the H37Rv reference genome (Rv2879c and Rv2880c) were targeted by their guide RNAs.^19^ Further, the fact that the frameshift mutation appears to be present in the H37Rv reference genome obscures the presence of *rlmN* in MTB by causing difficulty with annotation software; the current Mycobrowser reference genome does not list *rlmN* and the putative methyltransferase function of the gene is not listed under either Rv2879c nor Rv2880c.^15^ Recently, however, a comprehensive update to the H37Rv reference genome was released.^16^ While the reference genome still contains the *rlmN* frameshift mutation, improved annotation software has now recognized the homology of the gene fragment to *rlmN*. We propose that the community consider adopting this updated 2022 H37Rv reference genome in preference to the 1998 H37Rv reference genome with annotations from 2011 that is currently in use.

In conclusion, we report the novel association between a frameshift mutation in the methyltransferase *rlmN* and linezolid resistance in MTB, as measured by quantitative MIC. This statistical association is observed in a set of 7426 linezolid MIC phenotyped isolates. Further, we use protein structural studies to hypothesize a plausible causal mechanism of resistance from loss of function due to the ablation of the majority of the critical SAM cofactor binding sites.

As a part of this study, we provide three other items of broad usefulness to the scientific community: one, a catalogue of all known MTB methyltransferases, to be used as candidate genes for future drug resistance work for ribosomally-targeted drugs; two, a proposal that *rlmN* be considered as a candidate gene in future WHO mutation catalogues; and three, an evidence-based recommendation for use of the new, 2022 H37Rv MTB reference genome which benefits from updated annotations.

Our findings, which unveil a novel genetic mechanism underlying linezolid resistance in MTB, have future implications for improved resistance diagnostics and better-targeted therapeutic strategies.

## Online Methods

### Building a catalogue of confirmed and putative RNA methyltransferases in MTB

To find all confirmed and putative methyltransferases in MTB, we combined several strategies. First, we searched the standard 2011 H37Rv annotation furnished by Mycobrowser^15^ as well as a newly-released re-annotation of H37Rv^16^ for “methyltransferase” and “RNA”. We noted all genes and pseudogenes that were found in this manner (see Supplemental Information), whether or not they were included in the final list, and if not, why.

Additionally, we searched UniProt^14^ for methyltransferases thought to have RNA binding ability. Because some methyltransferases have dual specificity for tRNA and rRNA, we searched for any RNA methyltransferase that was either in the methyltransferase class (Enzyme Commission (EC) number 2.1.1.*),^31^ or that was annotated with the gene ontology (GO) term for RNA methyltransferase activity (GO: 0008173).^32,33^ Specifically, we used the following search terms:

Search 1: “ec:2.1.1.* H37Rv -H37Ra RNA”. We then looked for evidence of ribosomal methylation in the protein annotation.

Search 2: “GO:0008173 H37Rv -H37Ra”

Any protein that was found by these methods and that seemed likely to have RNA methyltransferase activity based on experimental evidence or homology was included in our catalogue.

### Confirmation that no additional *rlmN* homologs are present in MTB

To show that we did not miss any potential *rlmN* homolog(s) in MTB, we performed a tblastn search with the *S. aureus rlmN* protein sequence (UniProt ID: Q2FZ66) against the H37Rv reference genome and confirmed that the known MTB *rlmN* was the only hit produced. Default BLAST parameters were used.^17^

### Protein similarity analysis of RNA methyltransferases in MTB

Protein similarity analysis of the catalogue of confirmed and putative RNA methyltransferases in MTB (against the *S. aureus rlmN*) was performed using T-Coffee and the EMBL-EBI job dispatcher with the default parameters.^34,35^ Protein sequences were sourced from UniProt, using the accessions noted in the full text.^14^ The percent identity matrix was calculated with T-Coffee,^34^ and visualized using R version 4.4.2 and the pheatmap library.^36,37^

### Acquiring MTB isolate phenotype data from the BV-BRC

Information on MTB isolates with lab-based linezolid phenotypes was accessed from the BV-BRC on 31 July 2025.^20^ For analyses using binarized phenotypes, isolates were labeled resistant if their linezolid MIC was > 1.0 mg/L and sensitive otherwise, following the WHO recommended breakpoint.^22^ For all isolates with a lab-based phenotype value for linezolid, their BioSample Accession values were also accessed from the BV-BRC (n=8454).

### Processing uniformly-assembled genomes from AllTheBacteria

We used the BioSample Accession values from the BV-BRC^20^ to query AllTheBacteria,^21^ a resource supplying uniformly-assembled bacterial genomes. All genomes that were available were downloaded (n=8364). It was noted that a small number of genomes had unusual genome sizes (very large or almost empty) or otherwise poor assemblies. We proceeded to include for analysis only genomes with a size within 5% of the 4411532 bp size of the H37Rv reference genome (n=7775). We confirmed that genomes with aberrant sizes that were excluded were not enriched for resistant isolates (Before filtering: 1.71% resistance; after filtering, 1.79% resistance)

We then used BLAST to find our 4 genes of interest in each genome assembly.^17^ We used tblastn for the protein sequences (*rplC, tsnR, rlmN*) and blastn for the ribosomal DNA sequence (*rrl*). We only extracted the top hit for each search, with an E-value cutoff of 1e-10. After finding all sequences for each genome assembly, we only included for downstream analysis those isolates that had missense, nonsense, or frameshift/indel mutations (n=7436; 128 resistant, 7308 susceptible = 1.72% resistance), excluding those isolates with small, rare indels (present in <1% of either resistant or susceptible isolates). Again, the proportion of resistant isolates remained similar after filtering.

For each mutation detected in each gene, its presence or absence was noted in each isolate. We were then able to calculate the two-sided Mann Whitney U-test statistic for each mutation’s association with MIC in order to test the null hypothesis that isolates with and without a given mutation are selected from populations with the same MIC distribution. Resultant *p*-values were corrected using the Bonferroni procedure. Only mutations significantly associated with an increase in quantitative resistance (MIC) were reported in Table 2.

### Structural insights from *E. coli* RlmN

Protein sequence similarity between *rlmN* in MTB (UniProt ID: P9WH15) and in *E. coli* (UniProt ID: P36979) was performed with BLAST, using default parameters.^17^ The protein structure of RlmN was downloaded from the PDB (PDB ID: 3RFA).^39–41^ The protein-ligand interaction fingerprint was generated using Flare™ from Cresset®, and then modified for clarity and color-complementarity.^42^ Structural modelling was done in PyMOL version 3.1.3.^43^

## Declarations

### Ethics approval and consent to participate

Not applicable.

### Consent for publication

Not applicable.

### Availability of data and materials

Data supporting the conclusions of this article are available in the GitHub repository, https://github.com/BrynMarieR/rlmN_frameshift. Some data are also included within the article and its supplemental file. The other datasets mentioned in, but not created for, this study are available from their respective, publicly available repositories as mentioned in the main text.

### Authors’ contributions

B.M.R conceived and designed the study; acquired, analyzed, and interpreted the data; and drafted the manuscript. A.G.G. made substantial contributions to the conception and design of the study, aided in data interpretation, and contributed to the manuscript. All authors read and approved the final manuscript.

## Supporting information

Supplemental tables for genome annotation analysis

## Acknowledgements

The authors would like to acknowledge valuable discussions had with Professors Maha R. Farhat and M. Sloan Siegrist. The authors also thank members of the UMass SAGE lab for useful input. This work utilized resources from Unity, a collaborative, multi-institutional high-performance computing cluster managed by UMass Amherst Research Computing and Data.

